# Quantifying long-term predictability in microbial plankton dynamics

**DOI:** 10.1101/237743

**Authors:** Caterina R. Giner, Vanessa Balagué, Anders K. Krabberød, Isabel Ferrera, Albert Reñé, Esther Garcés, Josep M. Gasol, Ramiro Logares, Ramon Massana

**Affiliations:** Institut de Ciències del Mar (CSIC), Passeig Marítim de la Barceloneta, 37-49, ES-08003, Barcelona, Catalonia, Spain; University of Oslo, Department of Biosciences, Section for Genetics and Evolutionary Biology (Evogene), Blindernv. 31, N-0316 Oslo, Norway

**Author notes:** **Corresponding authors:**, Institute of Marine Sciences (ICM-CSIC) Passeig Marítim de la Barceloneta 37-49, 08003, Barcelona, Catalonia, Spain.

## Abstract

Determining predictability in community turnover is a key ecological question. In the microbial world, seasonality has been reported for communities inhabiting temperate zones, but not much is known on seasonality for individual species. Specifically, we have a vague understanding on the amount of species displaying predictability during temporal community turnover as well as on their dynamics. Here we developed a ‘*Recurrence Index*’ to quantify predictability in microbial species. Applying our index to 18S rDNA metabarcoding data from one of the longest temporal observatories of marine plankton we determined that 13% of the picoeukaryotic and 19% of the nanoeukaryotic species, accounting for about 40% of the community abundance in both fractions, feature predictable dynamics when sampled monthly during 10 years. Thus, most of the species analysed had unpredictable temporal abundance patterns. Altogether, we show that species with both predictable and unpredictable temporal dynamics can occur within the same seasonal microbial community.

## INTRODUCTION

A major challenge in ecology is to understand the mechanisms that determine community turnover across space and time. Community turnover can be explained by four main processes: selection, (ecological) drift, dispersal and speciation (Vellend 2010). Determining to what extent the combination of the latter four processes structure different communities is a current challenge for ecologists. This framework that derives mostly from the study of multicellular organisms has recently been applied to the microbial world (Hanson *et al.* 2012; Stegen *et al.* 2013). Microbes are key players in most ecosystems, yet we have a limited understanding of what governs their community turnover. For prokaryotes, environmental selection seems to be the most important process structuring communities (Lindström & Langenheder 2012) although there is also evidence indicating that stochastic processes (e.g. drift and dispersal) have a role (Ofiteru *et al.* 2010). Furthermore, multiple members of a community can be structured by distinct processes at different times, with some taxa responding, e.g., to selection, while others presenting stochastic assembly patterns (Langenheder & Szekely 2011; Stegen *et al.* 2013).

While many studies have concentrated on investigating spatial patterns of microbial community turnover (Lindström & Langenheder 2012), fewer have focused on exploring temporal dynamics (Fuhrman *et al.* 2015; Bunse & Pinhassi 2017). Given the relevance of microbial marine plankton for the functioning of the biosphere (Falkowski 2012), understanding the predictability in their dynamics should help comprehending their response to disturbance or global change. Ascertaining predictability in community turnover can also provide hints on the amount of functional redundancy present in the ecosystem (Allison & Martiny 2008), as communities dominated by functionally interchangeable taxa may exhibit a low degree of predictability in species composition with time. In the latter case, ecological drift could have a more relevant role than selection in determining community turnover (Chase 2003).

Annual cycles driven by meteorological seasons in temperate zones have clear effects on terrestrial ecosystems. In phytoplankton assemblages, yearly cycles in light, temperature and nutrients are known to induce clear abundance dynamics (estimated using chlorophyll-a concentration), generally with one or two abundance peaks per year, although cases with unclear patterns have also been observed (Winder & Cloern 2010). To date, different studies indicate that microbial community turnover is correlated with meteorological seasons (Fuhrman *et al.* 2015; Bunse & Pinhassi 2017), but most studies of microbial community dynamics have focused on prokaryotes, although eukaryotes also constitute key elements of microbial communities (Caron *et al.* 2009). One could expect that due to the structural and behavioural differences between eukaryotes and prokaryotes (Massana & Logares 2013; Keeling & Del Campo 2017) they may show different temporal dynamics. So far, temporal studies published (time series typically <5 years) hint at seasonality in microbial eukaryotic communities at the bulk level (Romari & Vaulot 2004; Countway *et al.* 2010; Kim *et al.* 2014; Genitsaris *et al.* 2015; Piredda *et al.* 2017).

Most studies on microbial dynamics have investigated whole community patterns, which are driven by abundant taxa. Investigating the temporal behaviour of individual taxa is important as some members of a microbial community could respond differently to cyclic environmental variation, while others may present no response. Focusing on individual taxa may also help detecting differences in abundance dynamics over time and in their persistence, with some species exhibiting i.e. smooth temporal fluctuations and others displaying rapid fluctuations. Furthermore, as most microbial communities are composed by a few abundant and a large number of low-abundant or rare taxa (Pedrós-Alió 2006, 2012; Logares *et al.* 2014; Logares *et al.* 2015), it is relevant to explore the temporal behaviour (i.e. the recurrence) across the taxa-abundance spectrum. Rare taxa can be metabolically active (Logares *et al.* 2015), respond to environmental change (Campbell *et al.* 2011; Lindh *et al.* 2015) and present repeatable community assembly patterns (Alonso-Saez *et al.* 2015). In temporal surveys, rare taxa can be assigned to one of three categories: (a) “Seasonal” taxa that were rare at the time of sampling, but which are systematically recruited to the abundant community during specific time periods, (b) “Opportunistic taxa” that are generally rare but become abundant exceptionally and for a short time [a.k.a. conditionally rare taxa, (Shade *et al.* 2014)], or (c) “permanently rare” taxa that never (within the limitations of the sampling design) become abundant (Logares *et al.* 2015).

Here we analyse and quantify the recurrence in the long-term community dynamics of microbial eukaryotes inhabiting an oligotrophic coastal site in the Mediterranean Sea [Blanes Bay Microbial Observatory, BBMO, (Gasol *et al.* 2016)]. Microbial eukaryotes were sampled monthly during 10 years in two size fractions and their diversity was assessed by high-throughput sequencing the 18S rDNA (V4 region). We developed a metric to detect and quantify temporal recurrence in different taxa (hereafter *Recurrence Index)*, which allowed us to determine that 13.2% and 18.6 *%* of the pico- and nanoeukaryotic Operational Taxonomic Units (OTUs), representing up to 40% of the relative abundance, featured predictable dynamics. We also observed that the overall system presents annual seasonality with two main recurrent configurations, and that these communities did not increase their differentiation along the 10 sampled years.

## MATERIALS AND METHODS

### Study site and sampling

We carried out a monthly sampling during 10 years at the Blanes Bay Microbial Observatory (BBMO) located in the North Western Mediterranean Sea (41°40’N, 2°48’E). This is a well-studied temperate oligotrophic coastal site that has relatively little human or riverine influence (Gasol *et al.* 2016). Surface water was sampled about 1 km offshore over a water column of 20 m depth, from January 2004 to December 2013. Water temperature and salinity were measured *in situ* with a CTD. Seawater was pre-filtered through a 200 μm nylon-mesh, transported to the laboratory under dim light in 20 L plastic carboys, and processed within 2 h. Samples for determination of chlorophyll-a concentration were filtered in GF/F filters, extracted with acetone and processed by fluorometry (Yentsch & Menzel 1963). Inorganic nutrients (NO_3_^−^, NO_2_^−^, NH_4_+, PO_4_^3-^, SiO_2_) were measured spectrophotometrically using an Alliance Evolution II autoanalyzer (Grasshoff *et al.* 1983). In statistical analyses, these variables were standardized as z-scores, that is, deviations of the values from the global mean.

About 6 L of the 200 μm prefiltered seawater were sequentially filtered using a peristaltic pump through a 20 μm nylon mesh, a 3 μm pore-size polycarbonate filter of 47 mm diameter (nanoplankton fraction, 3-20 μm), and a 0.2 μm pore-size Sterivex unit (Millipore, Durapore) [picoplankton fraction, 0.2–3 μm]. Sterivex units and the 3 μm filters were stored at -80°C. DNA extractions were performed at the end of the sampling period using a standard phenol-chloroform protocol (Schauer *et al.* 2003; Massana *et al.* 2004), with a final step of purification in Amicon units (Millipore). Nucleic acid extracts were quantified in a NanoDrop-1000 spectrophotometer (Thermo Scientific) and stored at -80°C until analysis.

### DNA sequencing and bioinformatics

The eukaryotic universal primers TAReukFWD1 and TAReukREV3 (Stoeck *et al.* 2010) were used to amplify the V4 region of the 18S rDNA (~380 bp). PCR amplification and amplicon sequencing was carried out at the Research and Testing Laboratory (<u>http://rtlgenomics.com/</u>) using the *Illumina* MiSeq platform (2x250 bp paired-end sequencing). *Illumina* reads were processed following an in-house pipeline (Logares 2017). Operational Taxonomic Units (OTUs) were delineated by clustering sequences at 99% similarity using UPARSE (Edgar 2013) as implemented in USearch v8.1. Only OTUs present in at least 3 samples were retained. Taxonomy was assigned roughly at class-level by BLASTing OTU representative sequences against PR^2^ (Guillou *et al.* 2013) and two in-house marine protist databases (available at https://github.com/ramalok) based in a collection of Sanger sequences (Pernice *et al.* 2013) or 454 reads (Massana *et al.* 2015). Metazoan, Streptophyta and nucleomorphs were removed. Two OTU tables were generated: (1) the pico-nano-eukaryotic OTU-table had 120 samples of picoeukaryotes and 89 of nanoeukaryotes (samples from May-2010 to July-2012 and from 4 additional dates were discarded due to suboptimal sequencing); (2) the picoeukaryotic OTU-table had 120 samples. To enable sample comparisons, both tables were randomly subsampled to the lowest number of reads per sample using the *rrarefy* function in *Vegan* (Oksanen *et al.* 2008). The pico-nano-eukaryotic table was subsampled to 5,898 reads per sample (14,771 OTUs), while the picoeukaryotic table was subsampled to 7,553 reads per sample (13,040 OTUs). The sequence data are publicly available at the European Nucleotide Archive with accession numbers XXXX.

### Recurrence analyses and community dynamics

We developed a *Recurrence Index* (RI) to identify taxa presenting temporal recurrence or predictable dynamics. To calculate the RI, we first applied the ACF (Auto-Correlation Function) comparing taxa (OTUs or taxonomic Classes) relative abundances at different time lags. Then, we summed the absolute ACF values for each taxon along the complete temporal series (RF). Afterwards, we compared the RF values against a null distribution obtained after randomizing 1,000 times the taxa abundances and calculating the absolute sums of the randomized ACF values. Subsequently, we calculated the mean of the null model (RF_random_) for each taxon plus its 97% confidence intervals (CI). The RI was calculated as: RI=RF/RF_random_. Based on empirical tests, a given taxon was considered recurrent if a) its RI was above a given threshold, here 1.20 for picoeukaryotes, and 1.15 for nanoeukaryotes, and if b) its RF was significantly higher than RF_random_ (that is, outside the CI and within the upper 1.5% probability). The code used to calculate RI is publicly available (Giner *et al.* 2017) and ready-to-use in R through EcolUtils (Salazar 2015). Detected recurrent picoeukaryotic taxa were classified to reflect how long was their presistance in the temporal-series according to their changes in abundance. For this, we identified for each taxon the number of months with abundances above the 10-year mean. Taxa displaying >30 months above this mean (i.e. at least 3 times per year, on average) were considered ‘long’-persistent, while those displaying less than 30 months were considered ‘short’-persistent.

To investigate recurrence in whole community dynamics, we computed the mean β-diversity (Bray-Curtis dissimilarities) for all pairs of samples taken *n* months apart (n ranging from 1 to 119 months for picoeukaryotes). In order to determine whether the observed β-diversity could be generated by random community dynamics, we calculated the Raup-Crick metric (Chase *et al.* 2011) using Bray-Curtis dissimilarities [hereafter RC_bray_] for the picoeukaryotes, following Stegen *et al.* (2013). RC_bray_ compares the measured β-diversity against the β-diversity that would be obtained under random community assembly. For each pair of communities, the randomization was run 999 times. Only OTUs with >500 reads were included in this analysis. RC_bray_ values > +0.95 or < -0.95 are interpreted as significant departures from a stochastic community assembly, pointing to deterministic assembly forces, such as environmental selection. On the contrary, RC_bray_ values between -0.95 and +0.95 point to stochastic community assembly (Chase *et al.* 2011).

### Community turnover and response of single OTUs to environmental variables

Non-metric multidimensional scaling (NMDS) was based on Bray-Curtis dissimilarity matrices. In NMDS, differences between predefined groups were tested with ANOSIM [ANalysis Of SIMilarity, (Clarke 1993)] performing 1,000 permutations. To determine the proportion of variation in community composition explained by environmental variables we used PERMANOVA. We also analysed the correlation between the matrix of environmental variables and that of community differentiation using Partial Mantel tests (Legendre & Legendre 1998) as well as by fitting environmental variables onto the ordination space of the NMDS *(envfit* function in *Vegan).* Finally, we performed an *IndVal* analysis [INDicator VALues, (Dufrene & Legendre 1997)] to identify OTUs associated to a specific season.

OTUs with statistically significant (p<0.05) *IndVal* values >0.3 were considered season-specific, following Logares *et al.* (2013). All analyses were performed using functions implemented in the packages *Vegan* (Oksanen *et al.* 2008), *pvclust* (Suzuki & Shimodaira 2006), and *Labdsv* (Roberts 2016) within the R Statistical environment (R-Development-Core-Team 2008).

Correlations between individual OTUs and environmental variables were done using extended local similarity analysis [eLSA] (Ruan *et al.* 2006; Xia *et al.* 2011). The analysis was based on the subsampled picoeukaryotic OTU-table together with the environmental variables. OTUs that were not present in at least 10 out of 120 months were excluded from the analysis, resulting in a dataset with 1,065 OTUs and 9 environmental variables. ELSA was run with default normalization (a z-score transformation using the median and median absolute deviation) and p-value estimations under a mixed model that performs a random permutation test of a co-occurrence only if the theoretical p-values for the comparison are <0.05. Missing data were interpolated linearly from adjacent months, and we did not allow any time delay.

Only the picoeukaryotic OTU table was used in rarity analyses. OTUs with abundances per sample that were always < 0.1% were considered permanently rare (Logares *et al.* 2015). To exclude the possibility that rare OTUs were aberrant variants of abundant ones, we only analysed rare OTUs that had a similarity <97% with any abundant counterpart. We considered as temporally abundant those OTUs with a mean abundance >0.1% along 10 years. Conditional Rare Taxa (opportunistic) were detected following the protocol described in Shade et al. (2014).

## RESULTS

### Quantifying recurrent patterns in different taxa

The *recurrence index* (RI) that we developed allowed to identify and quantify predictability in different taxonomic groups and OTUs. Overall, microbial eukaryotes present in the BBMO were very diverse and included more than 63 taxonomic groups at the class level, most of them found both in the pico- and nanoeukaryotic fractions, but with different relative abundances in the two fractions (Fig. S1). Picoeukaryotes were dominated by different alveolates (MALV-I, Dinoflagellata, MALV-II) and Mamiellophyceae (Fig. 1a), with the presence of many other groups at lower relative abundances, whereas nanoeukaryotes were dominated by Dinoflagellata and Diatomea (Fig. 2a). An inspection of the actual dynamics of the recurrent groups (RI >1.20 for picoeukaryotes, RI>1.15 for nanoeukaryotes) showed ACF values following cycles of 1-year periodicity, pointing to seasonality (see e.g. Mamiellophyceae in Fig. 1b). In picoeukaryotes, 13 of the 58 taxonomic groups tested exhibited a recurrent seasonal behaviour, and these accounted for 39.4% of the picoeukaryotic reads (Fig. 1a, Table S1). Two of the groups, MALV-III and Mamiellophyceae, exhibited a ‘strong-seasonal’ signal (RI>2.0), whereas the remaining 11 groups, including Dinoflagellata and some environmental clades, were ‘moderately-seasonal’ (1.2<RI<2.0). The remaining groups did not display recurrence. For the nanoeukaryotes, 13 groups exhibited seasonal behaviour (representing 8.2% of the nanoeukaryotic reads), and only MALV-III exhibited a ‘strong-seasonal’ signal (Fig. 2a, Table S2).

**Figure 1.**
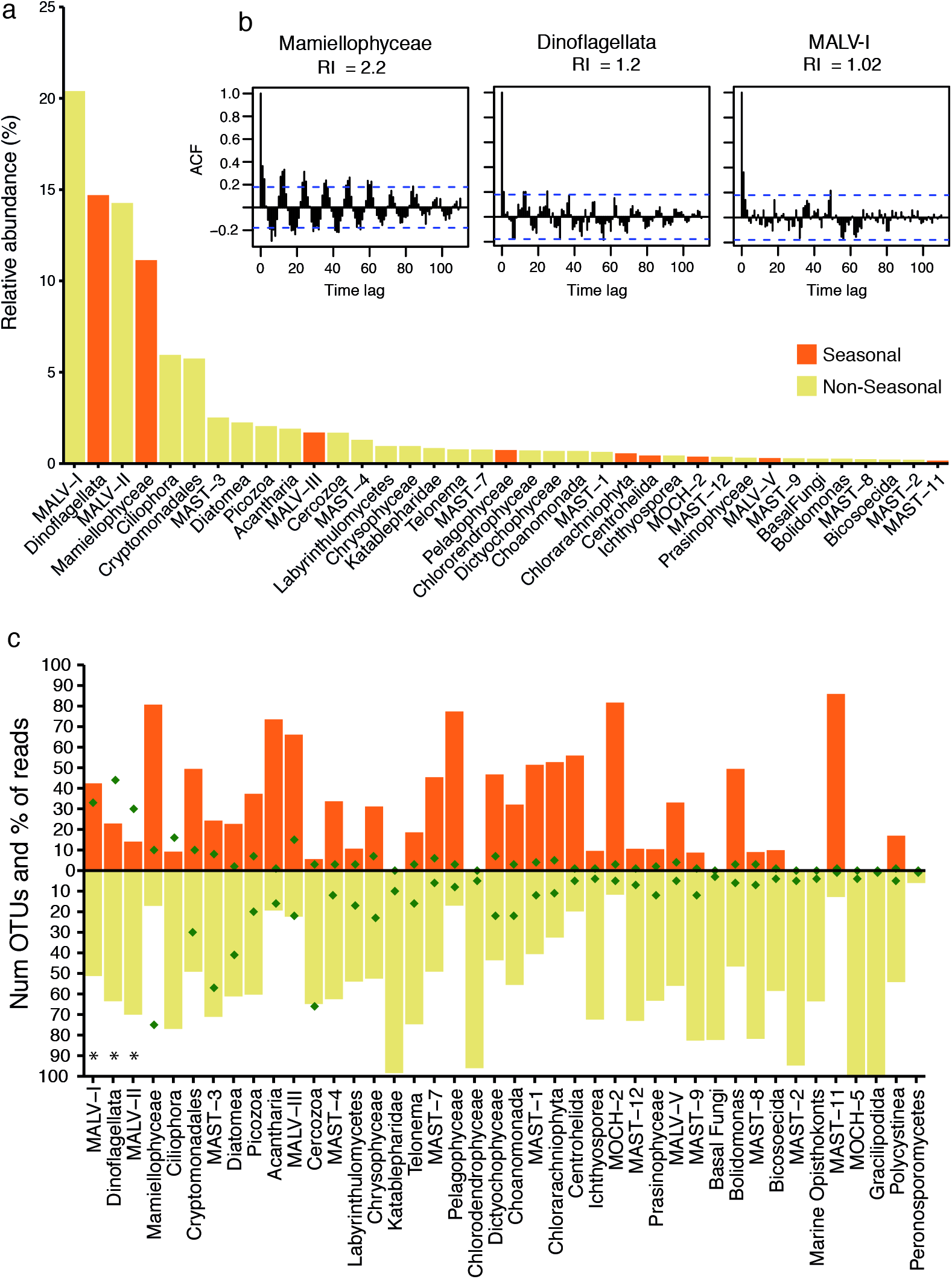
Taxonomic groups constituting the picoeukaryotic community in the BBMO and indications of their seasonality. (a) Average relative abundances of groups accounting for >0.2% of the reads. The bars are colored according to whether the group as a whole exhibits seasonality. (b) Selected autocorrelation function (ACF) plots showing ‘strong seasonality’ (Mamiellophyceae), ‘moderate seasonality’ (Dinoflagellata) and ‘no seasonality’ (MALV-I), together with their RI value. (c) Seasonal and non-seasonal signals for the main groups. The figure shows the number of OTUs (dots) and their percentage to all reads (bars) for each taxonomic group. Only OTUs present in >10 samples were considered.

**Figure 2.**
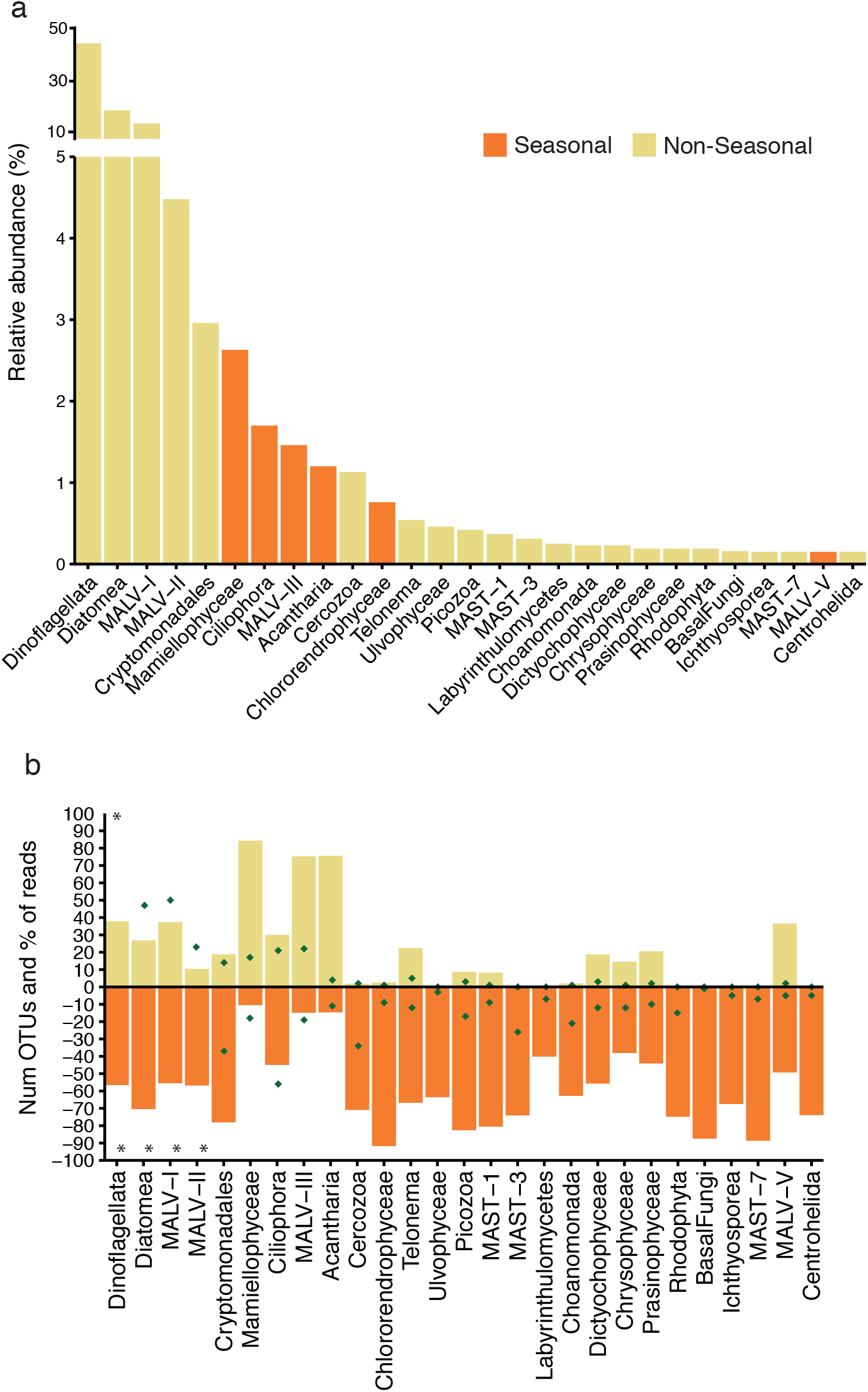
Taxonomic groups constituting the nanoeukaryotic community in the BBMO and indications of their seasonality. (a) Average relative abundances of groups accounting for >0.2% of the reads. The bars are colored according to whether the group as a whole exhibits seasonality. (b) Seasonal and non-seasonal signal for the main groups. The figure shows the number of OTUs (dots) and their percentage to all reads (bars) for each taxonomic group. Only OTUs present in >7 samples were considered. [* represents groups that have more than 100 OTUs].

We then explored the seasonal behaviour of individual OTUs, as seasonality at the taxonomic group level does not necessarily imply that all composing OTUs are also seasonal and, more interestingly, there can be seasonal OTUs in non-seasonal groups. We focused on the OTUs that were present in at least 10 samples for picoeukaryotes (1,898 OTUs representing ~90% of reads) and in at least 7 samples for nanoeukaryotes (2,266 OTUs representing ~91.5% of reads). Only 251 picoeukaryotic OTUs (representing 39.4% of the abundance) were seasonal. As expected, seasonal groups generally contained a majority of seasonal OTUs (Fig. 1c). Exceptions were low abundance groups (e.g. MALV-V, RAD-B) and the Dinoflagellata, which had a RI just above the cut-off (RI=1.23). We also identified recurrent OTUs in groups that did not show seasonality. In particular, Acantharia, Bolidomonas, Cryptomonadales, Dictyochophyceae, MAST-1, and MAST-10 had more reads (i.e. abundance) belonging to recurrent OTUs than to non-recurrent. Similar patterns were found in nanoeukaryotes (Fig. 2b), where 423 OTUs (accounting for 36.9% of the abundance) were seasonal.

Seasonal taxa exhibited different strategies based on how long was their persistence in the system (that is, ‘long’ and ‘short’ persistents; examples in Fig. S2). Nine out of the 13 seasonal picoeukaryotic groups (altogether accounting for 99.5% of the seasonal abundance) displayed long-persistence, while the remaining 4 groups (0.5% of the abundance) displayed short-persistence (Table S1). Persistence was also analysed for the 251 seasonal OTUs. About 31.5% (79 OTUs) showed long-persistence, belonging to groups that also displayed the same behaviour; these OTUs featured high relative abundances. The remaining seasonal OTUs had short-persistence. Furthermore, among the 89 rare OTUs that appeared in at least 10 samples, we detected nine that were ‘moderately-seasonal’ and displayed short-persistence.

### Community seasonality

To identify seasonality in community turnover, we calculated mean Bray-Curtis dissimilarities between samples separated by different time lags. Communities separated 12 months and their multiples (24, 36 and so on) showed the highest similarity, while those separated by 6 months and their multiples showed the highest dissimilarity for both pico- and nanoeukaryotes (Fig. 3). Thus, the investigated community displayed yearly seasonal patterns. Furthermore, community differentiation did not increase with time, as Bray-Curtis distances between samples separated by 1 year were very similar to those from samples separated by several years. Despite the observed dissimilarity cycling, community differentiation remained high during the 10 years, with averaged Bray-Curtis values ranging between 0.7 and 0.9.

**Figure 3.**
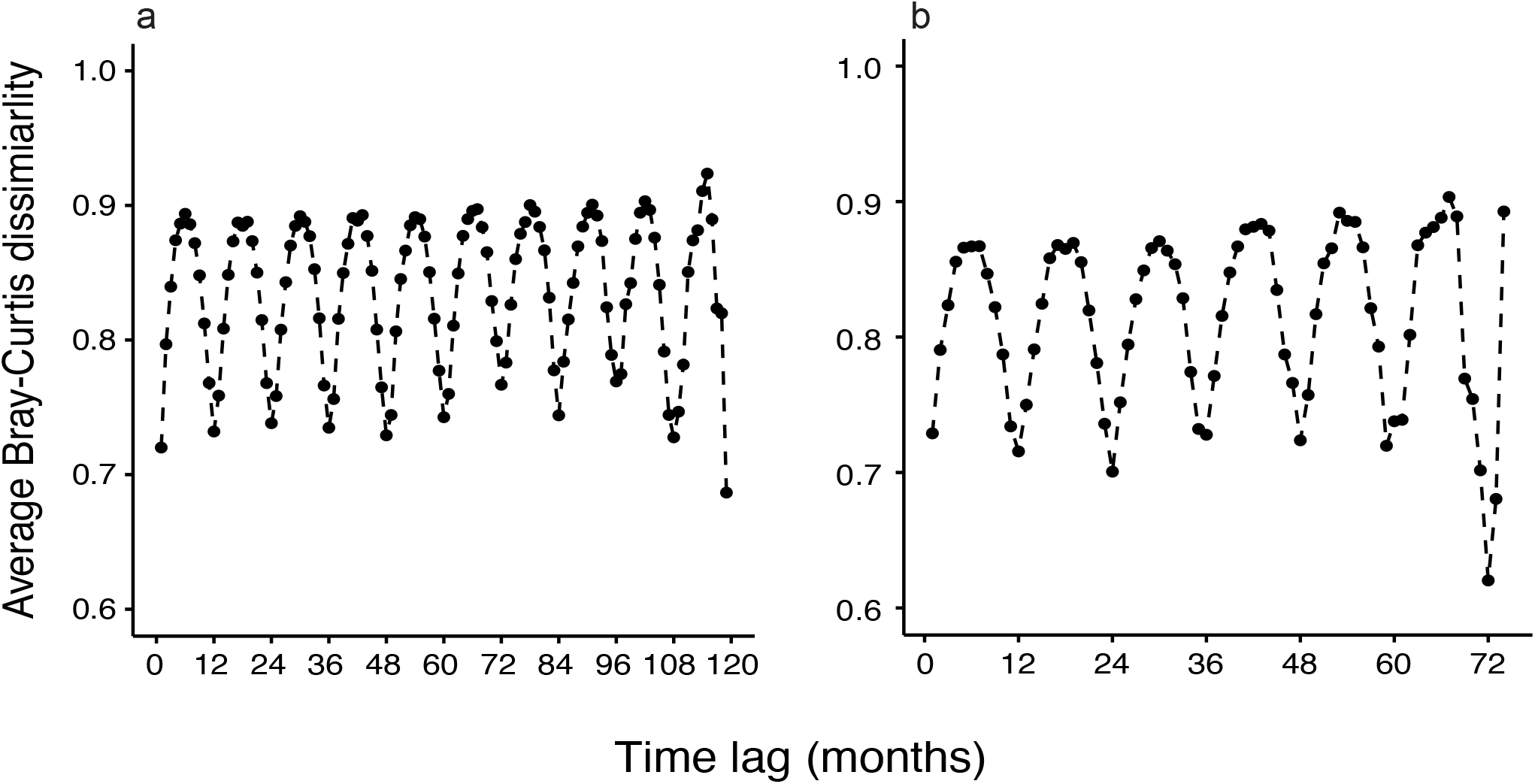
Inter-annual recurrence of communities of picoeukaryotes (a) and nanoeukaryotes (b), shown by the average Bray-Curtis dissimilarities of all pairs of communities separated by a given number of months (from 1 to 119 in a) and from 1 to 74 in b)).

Further analysis of whole community turnover unveiled two main recurrent structuring patterns throughout the 10 years. First, picoeukaryotes and nanoeukaryotes formed different groups characterized by their different cell sizes (Fig. S3). Second, samples from both assemblages formed two clearly marked groups corresponding to winter and summer months (ANOSIM test: R_pico_=0.717; R_nano_=0.713, p<0.001; Table S3). In contrast, spring and autumn communities did not form defined groups. In addition, winter communities were more similar among themselves when compared to other intra-seasonal variability (Fig. S4). For picoeukaryotes, we detected 173 season-specific OTUs, most of them associated to the winter and summer states (56 and 59 OTUs respectively, Table S4).

The seasonal patterns at the whole community level shown above were driven by the most abundant OTUs, which have a stronger weight in Bray-Curtis dissimilarities. Therefore, it was relevant to investigate whether the rare biosphere exhibited any seasonality. Within picoeukaryotes, 3,095 OTUs were considered permanently rare. Similar to what we found for the entire community, we observed two main rare community states associated to winter and summer months (Fig. S5a) with spring and autumn communities being transient states. We also found that the averaged Bray-Curtis values were most similar between rare communities separated by 1 year (and their multiples), and most different when separated by half a year (Fig. S5b). Bray-Curtis values considering rare taxa were higher than the ones for the entire community (ranging from 0.9 to almost 1), indicating that despite clear evidence of seasonality, the rare sub-community was very different from year to year.

### Environmental factors and species turnover

The BBMO site featured annual cyclic fluctuations of environmental conditions: the days were longer in early summer, water temperature was maximal two months later, and inorganic nutrients, particularly nitrate, nitrite and silicate, peaked in winter (Fig. S6). Selected environmental variables were fitted to the NMDS separately for picoeukaryotes and nanoeukaryotes (Fig. 4). In both cases, day length and temperature were the variables better correlated with community turnover (envfit; day length: r^2^=0.62, temperature: r^2^=0.56, p<0.001; Table S5). When controlled by each other in partial mantel tests, both variables still presented a moderate significant correlation with community composition (r=0.44 temperature, r=0.40 day length, p=0.001). The remaining environmental variables presented weaker or non-significant correlations with community composition (Table S5). Additional analyses indicated that a large part of community variance (76.8% in PERMANOVA) was not explained by any of the measured environmental variables. Day length and temperature, together, explained only 16% of community variance (p<0.001) in PERMANOVA analysis, a value that increased to 26% when running the analysis only with the OTUs that showed recurrence. Even though environmental variables explained a minor part of community turnover, results from the RC_bray_ analyses indicated that 93.4% of the measured β-diversity differed from what it would be observed under random community assembly, thus suggesting environmental selection. Specifically, 86% of the β-diversity comparisons presented RC_bray_ > +0.95, and 7.4% RC_bray_ < -0.95. Furthermore, by using eLSA analyses we detected 2,375 OTUs that were positively or negatively correlated with the analysed environmental variables (Table S6); these OTUs tended to be abundant. Specifically, 4% of the OTUs, representing ~47% of the total abundance, were positively or negatively correlated with temperature or day length.

**Figure 4.**
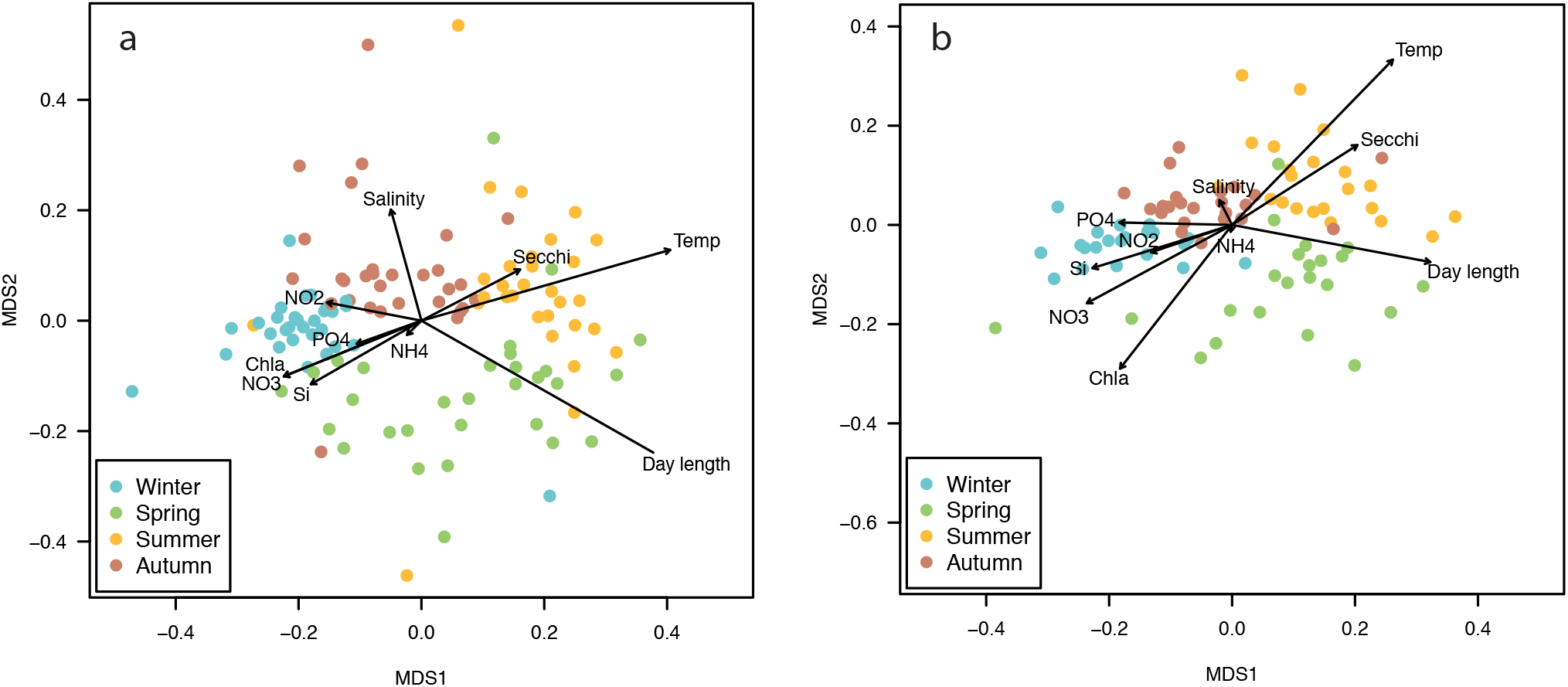
NMDS analysis of protist communities in monthly samples taken during 10 years in the BBMO showing the environmental vectors that better fit the plot after an Envfit test for the picoeukaryotes (a), and the nanoeukaryotes (b).

### Diversity patterns

Most individual samples (~80%) were close to richness saturation (details not shown). We also found richness saturation when constructing rarefaction curves based on the entire dataset of pico- and nanoeukaryotes (Fig. S7a), indicating that we recovered most of their diversity present in the BBMO throughout the 10 years. In accumulation curves, richness increased rapidly until approximately the 60^th^ month of sampling, and subsequent samples contributed little to new OTUs (Fig. S7b). Alpha diversity presented clear temporal trends.

For both size fractions, averaged richness and Shannon indices were highest during the autumn and winter months and significantly lower during spring (Fig. 5, p<0.05 Wilcoxon test). No statistical differences were found between pico- and nanoeukaryotes when comparing all samples together, nor when we compared each of the seasons separately (Wilcoxon tests, p>0.05).

**Figure 5.**
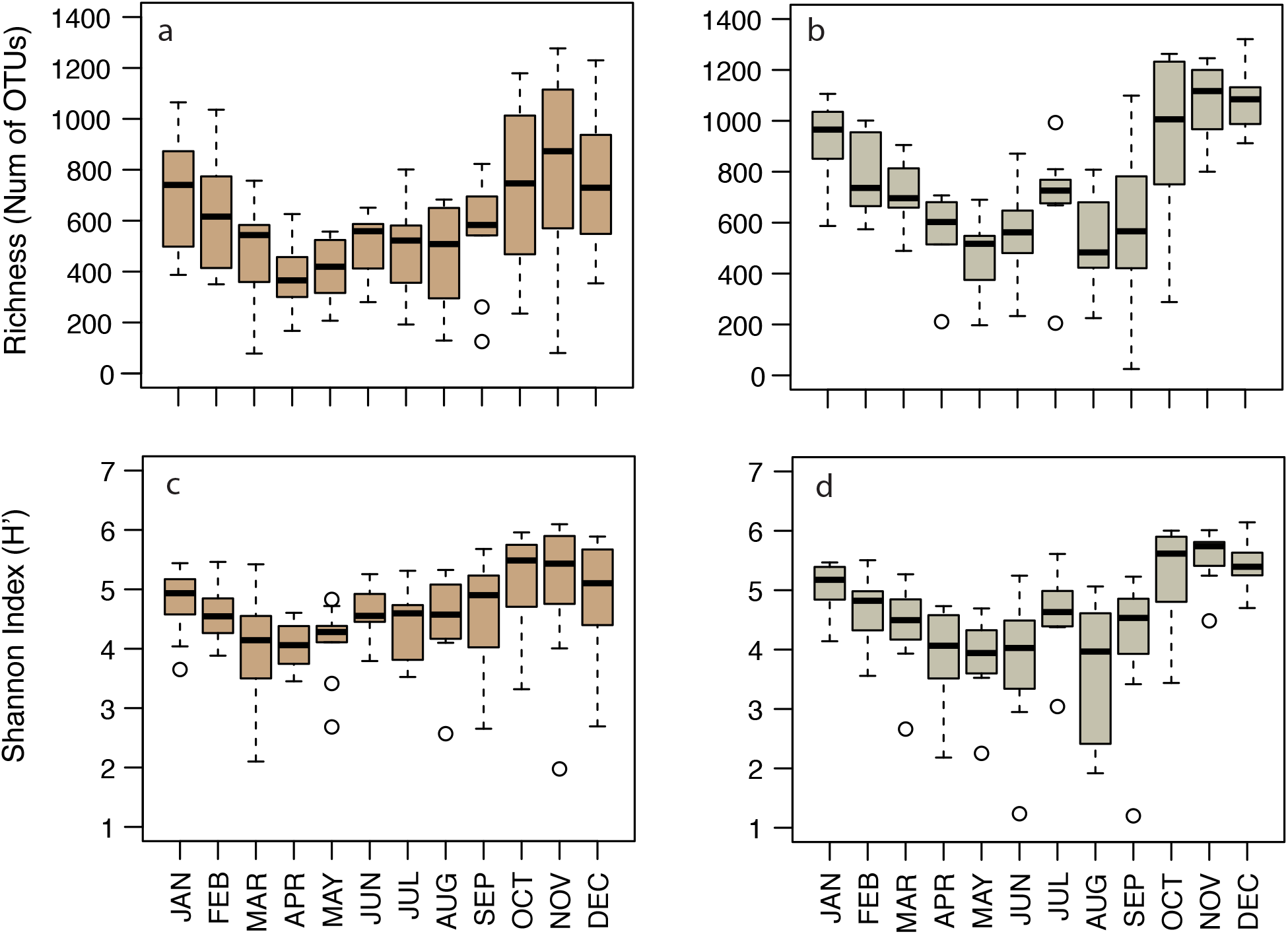
Monthly variation of alpha diversity present in BBMO protist communities. Boxplots display the monthly variability during 10 years of the richness (a, b) and Shannon indices (c, d) for the picoeukaryotes (a, c) and nanoeukaryotes (b, d).

## DISCUSSION

Predictable patterns in microbial long-term seasonal dynamics have previously been reported (Fuhrman *et al.* 2015; Bunse & Pinhassi 2017). Yet, previous studies have provided limited information on the proportions of the community that display predictable vs. unpredictable dynamics; see e.g. Winder and Cloern (2010). Here, using data from one of the longest protistan time-series analysed to date, we quantified repeatability in species and group seasonal dynamics along 10 years using a newly developed *Recurrence Index* (RI). We found that the microbial eukaryotic plankton includes species showing predictable as well as unpredictable temporal patterns.

### Quantifying community seasonality and predictability

Until now, one of the limitations in studies of microbial seasonality was the lack of routines to quantify recurrence patterns in different taxa. Our *recurrence index* (RI) allowed us to quantify the recurrence of all taxonomic groups and OTUs in the analysed temporal series. We found seasonality in only 13.2% and 18.7% of the pico- and nanoeukaryotic OTUs respectively (yet accounting for 39.4% and 36.9% of the abundance). Using the RI, we also identified seasonality at taxonomic group levels. In particular, Mamiellophyceae and MALV-III featured strong seasonality. As expected, most OTUs within these groups were also seasonal, indicating that this was a conserved trait in most species of the group. The strong seasonality of Mamiellophyceae was already suggested, as species of this group seem to have a preference for low temperatures (Foulon *et al.* 2008), whereas the strong seasonality of MALV-III is intriguing, given that virtually nothing is known for this group. The opposite scenario, seasonal OTUs within non-seasonal groups, was also found and was explained by contrasting dynamics in different OTUs within a group, MALV-II being a good example. In most cases, seasonal groups and OTUs displayed one peak of abundance per year, despite our

RI can detect other patterns of abundance dynamics (i.e two peaks of abundance per year, one peak every two years and so on). In a study using Chlorophyll-*a* as a proxy of abundance in 125 time series studies, Winder and Cloern (2010) found that having one annual abundance peak is the most common pattern among phytoplankton (detected in ~48% of the analysed time series).

Our results demonstrated that pico- and nanoeukaryotic communities displayed yearly seasonality throughout the 10 years. Furthermore, as shown by the Raup-Crick analyses, we found that β-diversity was significantly different from chance, indicating that community dynamics were not driven by ecological drift (Chase *et al.* 2011; Stegen *et al.* 2013). Interestingly, there was no temporal trend in community dissimilarity, as the difference between communities did not increase with time along the 10 years. This suggests seasonal recurrence of at least some abundant taxa and absence of mass effects [that is, massive immigration] (Lindström & Langenheder 2012). A somewhat contrasting result was found by Chow et al. (2013) in surface marine prokaryotic communities sampled monthly during 10 years, as dissimilarity increased slightly during the first fourth years. In our study, averaged Bray-Curtis dissimilarity values between any pair of samples was ~0.8, while in Chow et al. (2013) was ~0.6, indicating that seasonal communities are far from identical during their turnover, and that there is not a unique community state to return, leaving room for multiple community configurations that likely reflect historical processes or ecological drift (Chase 2003).

Overall, the repeatability in community turnover points to a certain degree of resilience and predictability in the system (Fuhrman *et al.* 2015), although not all species may show these patterns. We found that the majority of OTUs did not present recurrence patterns or predictability (or it could not be detected due to the low signal), but the few that did were particularly abundant. This suggests that the community includes certain amounts of functional redundancy (Allison & Martiny 2008) by which different but ecologically similar OTUs become dominant in different years. Nevertheless, the link between richness and ecosystem function is still unclear. There is evidence indicating that changes in microbial richness affects ecosystem processes (Allison & Martiny 2008; Peter *et al.* 2011), thus pointing towards limited functional redundancy, as well as evidence supporting redundancy (Lyons & Dobbs 2012). Further studies are needed to determine the role of ecological redundancy in microbial plankton dynamics.

Seasonal taxa presented different persistence times, which could point to different strategies according on the time spanned between samplings. Short-persistents might reflect a faster growth under the presence of specific resources and a faster decrease perhaps due to a high predation, competitive pressure or viral mortality. On the other hand, long-persistents could reflect relatively slow growth (and slow use of resources) accompanied with relatively low predation or competition pressures, thus maintaining taxa in the system for relatively longer periods. Long-persistents may also have their growth rate tightly associated to some environmental variables (e.g. temperature), with their abundance reflecting environmental variability. We have also found that 1.6% of the OTUs were Conditionally Rare Taxa (CRT), a magnitude coinciding with that observed by Shade and Gilbert (2015) for prokaryotes. We considered these OTUs opportunistic, with an increase in abundance triggered by environmental cues. Finally, 23.7% of the OTUs were permanently rare, and only 9 of them showed seasonality, similarly to what was observed in marine bacterioplankton (Alonso-Saez *et al.* 2015). The permanently rare sub-community mirrored the recurrent annual pattern found in the whole community, showing that processes driving community turnover act on abundant as well as on rare species. Yet, the overall dissimilarity values among samples from the rare sub-community were higher than the values observed for the whole community, implying a larger stochasticity in the rare sub-community dynamics. However, detection limits for rare taxa may have inflated β-diversity estimates (Leray & Knowlton 2017). Interestingly, the latter supports the idea that communities include rare species which are metabolically active, as suggested by their seasonality, but never become abundant as they are adapted to a low abundance life (Logares *et al.* 2015) or subjected to high mortality pressures.

### Community and OTU response to environmental variation

Similar to bacterioplankton (Fuhrman *et al.* 2015; Bunse & Pinhassi 2017), we hypothesized that environmental selection would be a major force driving protist community seasonality. However, the measured environmental factors explained a minor fraction of community variability along the 10 years, which agrees with a previous study of protists showing that environmental fluctuations were explaining ~30% of community seasonality (Genitsaris *et al.* 2015). A possible explanation is that environmental selection has different effects on microbial eukaryotic and prokaryotic plankton (Logares *et al.* 2017), thus affecting their dynamics. Nevertheless, the studied system oscillated between two main community configurations that corresponded to winter and summer months, pointing to some degree of environmental selection. Interestingly, we found that a small fraction of pico-eukaryotic OTUs (~4%), yet representing ~47% of the total abundance, correlated positively or negatively with temperature and day length; temperature is an important factor structuring marine bacterioplankton communities across space (Sunagawa *et al.* 2015) and time (Fuhrman *et al.* 2006; Chow *et al.* 2013). Thus, we found evidence of environmental selection associated to variables that contrast between summer and winter, but such selection seems to be experienced by only a subset of the taxa in the community. Overall, differential or limited responses of OTUs to environmental variation may explain the low correlation between whole community and environmental variation, as contrasting OTU dynamics may cancel out or generate noise in whole-community analyses. Furthermore, the presence of two community states suggests that environmental selection intensity may change throughout the year, being stronger in summer and winter, and weaker in spring and autumn, thus allowing for ecological drift (Chase 2003) in the latter seasons. This is consistent with the larger number of OTUs exclusively associated to summer and winter months as compared to those associated to autumn and spring. In addition, the fact that winter communities were the most similar along the 10 years indicate a stronger environmental selection during this period compared to summer. Besides, autumn and spring may be intrinsically more variable seasons, with episodic rains and less constant temperatures/irradiances, representing year-specific environmental selection regimes. In particular, spring and autumn communities appeared as transitional states. The existence of two main states and two transitional states has previously been reported for Atlantic Ocean bacterioplankton (Ward *et al.* 2017), while the analysis of protist dynamics during two and a half years in the English Channel revealed three seasonal states corresponding to summer, autumn-winter, and spring (Genitsaris *et al.* 2015). Overall, the existence of recurrent states associated to different seasons suggests that environmental selection drives, to certain extent, community dynamics. Finally, ecological interactions can also have a role in community turnover, yet it has been suggested that they affect dynamics in periods ranging from days to weeks (Bunse & Pinhassi 2017).

In summary, by applying our *recurrence index* to data from one of the longest microbial time series to date, we show that microbial plankton communities include both taxa with predictable as well as unpredictable dynamics. Our quantifications indicate that 13.2% and 18.6% of the pico- and nanoeukaryotic OTUs representing 39.4% and 36.9% of the abundance respectively were seasonal or predictable. Thus, most taxa in our system had unpredictable temporal dynamics or we could not detect it. To our knowledge, this is the first time that both behaviours are reported to coexist in microbial plankton communities and quantified at the OTU level. Future studies need to determine whether the latter amount of seasonal predictability is typical of microbial communities in other temperate zones.

## ACKNOWLEDGEMENTS

We thank all members of the Blanes Bay Microbial Observatory sampling team and the multiple projects funding this collaborative effort over the years. The data and analyses done here have been funded by the Spanish projects FLAME (CGL2010-16304, MICINN) and ALLFLAGS (CTM2016-75083-R, MINECO) to RM and INTERACTOMICS (CTM2015-69936-P, MINECO/FEDER, EU) to RL and the European project DEVOTES (grant agreement no. 308392) to EG. CRG was supported by a FPI fellowship. RL was supported by a Ramón y Cajal fellowship (RYC-2013-12554, MINECO, Spain). We thank Guillem Salazar for his help with R analyses. Bioinformatic analyses have been performed at the Marbits platform (ICM-CSIC; <u>https://marbits.icm.csic.es</u>).

